# Object visibility, not energy expenditure, accounts for spatial biases in human grasp selection

**DOI:** 10.1101/476101

**Authors:** Guido Maiello, Vivian C. Paulun, Lina K. Klein, Roland W. Fleming

**Affiliations:** Department of Experimental Psychology, Justus-Liebig University Giessen, Giessen 35394, Germany

**Keywords:** Precision grip, Movement distance, Minimum energy, Object visibility, Perception/action, Reaching/grasping, Visuo-haptic interactions

## Abstract

Humans exhibit spatial biases when grasping objects. These biases may be due to actors attempting to shorten their reaching movements and therefore minimize energy expenditures. An alternative explanation could be that they arise from actors attempting to minimize the portion of a grasped object occluded from view by the hand. We re-analyze data from a recent study, in which a key condition decouples these two competing hypotheses. The analysis reveals that object visibility, not energy expenditure, most likely accounts for spatial biases observed in human grasping.

## Main Text

Human grasp selection is influenced by an array of factors, including the size, shape, mass, material, orientation, and position of the grasped object (e.g. see Cesari & Newell, 1999; Paulignan, Frak, Toni, & Jeannerod, 1997; Paulun, Gegenfurtner, Goodale, & Fleming, 2016; Schot, Brenner, & Smeets, 2010). Additionally, it has been proposed that humans may attempt to perform grasping movements economically, i.e., by minimizing the amount of work and resulting energy expenditure (Huang, Kram, & Ahmed, 2012). Minimizing energy expenditures could therefore explain spatial biases in grasping patterns, such as the biases toward shorter movement distances observed in several studies (Desanghere & Marotta, 2015; Glowania, van Dam, Brenner, & Plaisier, 2017; Kleinholdermann, Franz, & Gegenfurtner, 2013). However, a study by Paulun, Kleinholdermann, Gegenfurtner, Smeets, & Brenner (2014) questions this hypothesis. Participants were asked to grasp objects while approaching them from different sides. Contrary to the expectation that participants should be biased toward shorter reaching movements regardless of the side of approach, the authors found that participants grasped the right side of the objects irrespective of where the movement started when grasping with the right hand. The authors concluded that participants simply preferred grasping objects on the side of the acting hand, and suggested that this behavior may help increase the visibility of the objects during grasping and subsequent manipulation (Bozzacchi, Brenner, Smeets, Volcic, & Domini, 2018).

A more recent study by Paulun et al. (2016), which investigated how material properties and object orientation affect grasping, serendipitously contained two experimental conditions that can be used to contrast the object visibility hypothesis against the minimum reach hypothesis (Figure 1). Participants were asked to grasp, with a precision grip, small cylinders of Styrofoam, beech wood, brass and Vaseline-covered brass presented at different orientations. In the 150-degree rotation condition (Figure 1a), grasping the object on its right side would result in shorter reach movements as well as increased object visibility, whereas grasping the object on its left side would result in longer reach movements as well as decreased object visibility: here the object visibility and minimum reach hypotheses make positively correlated predictions. The two hypotheses make inversely correlated predictions in the 60-degree rotation condition (Figure 1b). Here, grasping the object on its right side would result in longer reach movements but increased object visibility, whereas grasping the object on its left side would result in shorter reach movements but decreased object visibility.

**Figure 1.**
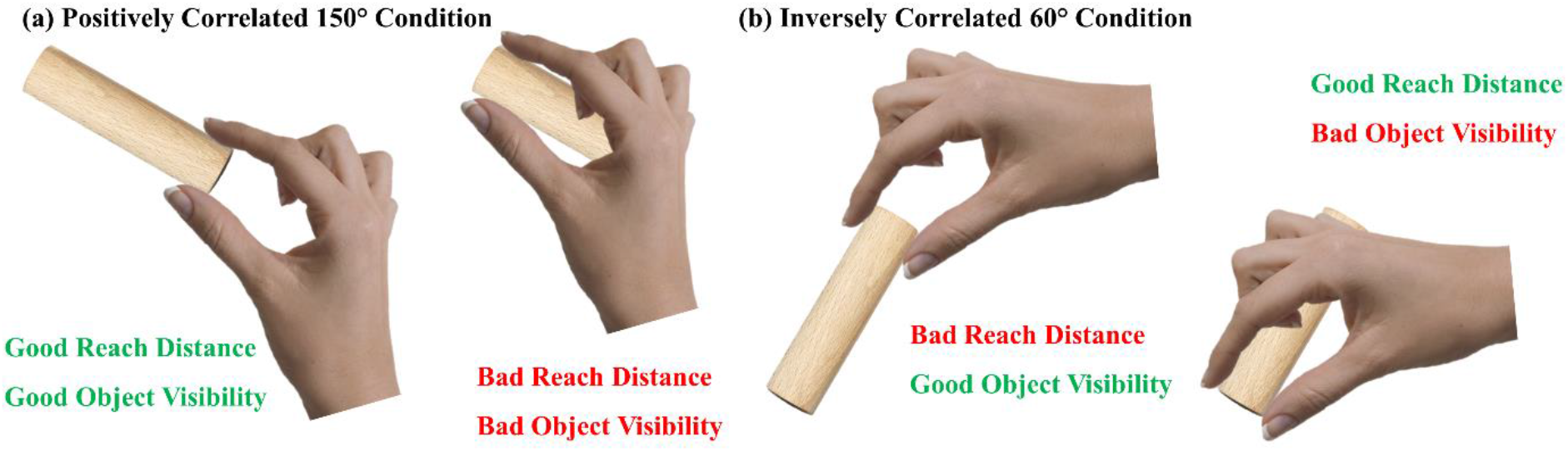
Two conditions from Paulun et al. (2016) that contrast the object visibility and minimum reach hypotheses against each other.

We therefore reanalyzed the data from these two conditions from Paulun et al. (2016) to distinguish whether participants (N=14) exhibited grasping behavior consistent with the minimum reach or the object visibility hypotheses. In Paulun et al. (2016) participants sat in front of a table to perform the grasping movements. Targets were placed in front of the participants, 36 cm away from the table edge. In each trial, participants positioned their right hand at a start location 11 cm away from the table edge and 26 cm to the right of the object (thus 36 cm from the object center). Following an auditory cue, participants grasped the stimulus object with a precision grip, lifted it, and transported it to a goal position of 13 cm diameter located 28.5 cm to the right of the object (center to center) and elevated 3.7 cm from the table. The position of the tips of the thumb and index finger were recorded using an Optotrak 3020, the position of the fingertips at the moment of first contact were determined using the methods adapted by Schot, Brenner, & Smeets (2010b). Participants executed 5 trial repetitions for each condition (4 materials x 6 orientations; here we only consider the 150-deg and 60-deg orientation conditions).

In our reanalysis for both the 150-deg and 60-deg conditions, we first computed the medoid grasp for each participant across object materials and trial repetitions (i.e. 20 trials per observers and orientation), and then we computed the medoid grasp across participants. The medoid (a concept similar to the mean) is the element of a set that minimizes its distance to all other elements. We excluded from the analysis the 4% of grasps that fell along the long axis of the objects. First, we looked at the medoid grasp pattern in the 150-degree rotation condition and confirmed that the medoid grasp across participants was biased to the right side of the object (Figure 2a). We quantified the bias as the mean deviation of the grasp center (average between thumb and index finger) from the object midline. Next, we used the bias in the 150-deg condition to make predictions regarding what the bias should be in the 60-deg condition under the two competing hypotheses. Additionally, we made the simplifying assumptions that grasps should be perpendicular to and in contact with the surface of the object. Thus, in the 60-deg condition, if participants were attempting to increase object visibility, they should exhibit a similarly-sized bias for grasps above the object midline (Figure 2b). If, on the other hand, participants were attempting to minimize reach distance (and therefore energy expenditures), grasps should be biased by the same amount to the region below the object midline (Figure 2b). Figure 2d shows how the medoid grasp across participants and conditions is indeed shifted above the object midline, contrary to the minimum reach hypothesis, and in near perfect alignment with the object visibility hypothesis. This observation is confirmed by a simple statistical test: the average grasp distance to the object visibility prediction, across participants, is significantly smaller than the average grasp distance to the minimum reach prediction (t(13)=5.66, p=7.8*10^−5^, paired samples t-test).

**Figure 2.**
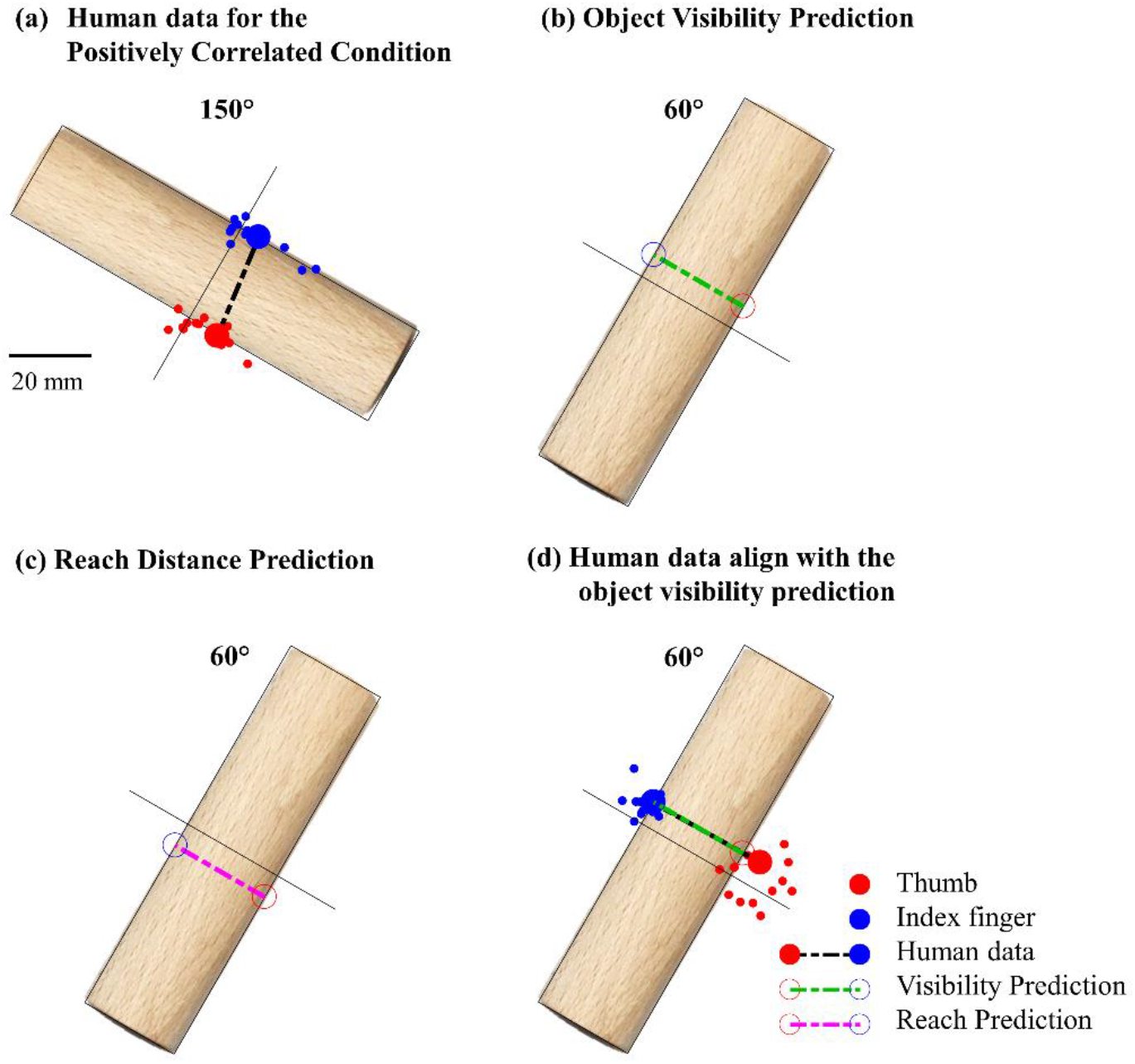
Human grasps compared to the two competing hypotheses. Small markers represent human medoid grasps for each participant across object materials and trial repetitions. Large markers are the medoid grasp across participants.

Our observation therefore suggests that humans are not attempting to minimize energy expenditures when selecting where to grasp an object, at least not through minimizing reach distance. Instead, the observed spatial biases for which participants tend to grasp objects on the side of the acting hand are consistent with the hypothesis that humans are attempting to minimize the portions of the objects occluded by the hand. Energy minimization principles may still play a role in the planning and on-line control of arm and hand movements during grasping (e.g. Soechting, Buneo, Herrmann, & Flanders, 1995). However, in the situations in which spatial biases in grasping are typically observed (Desanghere & Marotta, 2015; Glowania et al., 2017; Kleinholdermann et al., 2013), these biases are likely too small to induce noticeably different energy costs. Therefore, object visibility, not energy expenditure, accounts for these spatial biases in human grasp selection.

## Data availability

Data and analysis scripts are available from the Zenodo database (doi:10.5281/zenodo.2247283).

## Acknowledgments

This research was supported by the DFG (IRTG-1901: “The Brain in Action” and SFB-TRR-135: “Cardinal Mechanisms of Perception”), and an ERC Consolidator Award (ERC-2015-CoG-682859: “SHAPE”). Guido Maiello was supported by a Marie-Skłodowska-Curie Actions Individual Fellowship (H2020-MSCA-IF-2017: “VisualGrasping” Project ID: 793660).

Author Contributions
GM, VCP, LKK and RWF conceived and designed the study. VP collected the data. GM and VP analyzed the data. All authors wrote the manuscript.

